# Emergence of neural persistence: Insights from computational modelling

**DOI:** 10.1101/2024.02.19.581018

**Authors:** Vishal Verma

**Keywords:** Excitation Inhibition ratio, Wakefulness, Phase Transitions, Neural Network, Nonlinear Dynamics, Percolation, Self-organization, ordered-disordered Systems, Gamma Oscillations, Spontaneous Brain Activity, Symmetry and Symmetry Breaking

## Abstract

The persistent neural activity at a global scale, either stationary or oscillatory, can be explained by the use of the excitatory-inhibitory neural network models. This state of the network, as can be inferred, is crucial for the information processing and the memorizing ability of the brain. Though the goal for persistence to exist is apparent; from where the network achieves its ability to show a rich variety of the persistent dynamical states is unclear. The following study investigates the possible reasons for the persistence of neuronal networks in two parts; numerically and analytically. Presently, it shows that the action of the inhibitory components, among other favourable factors, plays a key role in starting and stabilizing neural activity. The results strongly support previous research conducted with both simpler and more specialized neural network models, as well as neurophysiological experiments.

**PACS numbers (2006 scheme):** 05.40.-a, 05.45.-a, 87.00, 89.00

## 1 Introduction

The concept of permanence (or persistence) was first introduced in applied mathematics during the 1980s for the study of three interacting predator-prey populations and then applied to the host-parasite populations, see, e.g., [1]. Various definitions of the term regarding such systems from time to time were given until, Butler *et al* [2] clearly defined the concept of persistence for the first time and classified it as weak-uniform and uniform persistence. They laid out the conditions under which the weak persistence of a dynamical system (with respect to the boundary conditions of the given set) does imply uniform persistence. They defined persistence as follows. Let *x*_1_, *x*_2_, *x*_3_, …, *x*_*n*_ be the dynamic variables of a model ecological/physical/biological system. The model was defined by the equation (1) with an imposed constraint.

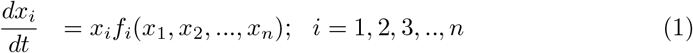

The constraint that *x*_*i*_ ≥ 0, i.e., all solutions must be nonnegative for all times must be satisfied. This constraint shall cause the solutions to remain bounded within a nonnegative cone in space *R*^*n*^. The boundary of the cone is a barrier for all the solutions for the dynamical system. If this boundary is invariant, then the solutions that are zero at any time shall remain so for all times. To allow for the dynamical system under the question to exist persistently, nonnegative solutions must remain away from the boundary of the cone at all times, to preempt the possibility of them sliding to zero. Generally speaking, the term persistence is given to systems in which strictly positive solutions *do not approach* the boundary of the nonnegative cone as *t* → ∞. When it is required only that *strictly positive* solutions do not asymptotically approach the boundary as *t* → ∞, the resulting system is said to be weakly uniformly persistent. When the system follows the rule that each strictly positive solution is eventually at some positive distance from the boundary; it is uniformly persistent. The latter statement implies that the strictly positive solutions are eventually uniformly kept away from the boundary. The classification was furthered by Freedman, one of the authors, with Moson in 1990 [3]. This definition can be understood in the context of the neuron if its resting potential value (nearly -75 mV) is considered at makeshift zero. All this was an analytical enterprise. My paper numerically analyses models for the neural networks of the brain to ask significant questions pertinent to brain dynamics, viz., (1) What parameters shall be responsible for bounding the neural voltages in the network away from the boundary and causing permanence? (5) What can be the simplest dynamical system that generates complex sustained dynamics while also shedding light on the mechanism by which such dynamics is produced in the brain? The answer to the former query is as follows: Since a network consisting of excitatory and inhibitory (EI) neurons is considered a canonical model for understanding cortical network activity, I will attempt to answer the rest of the questions using this model while also invoking a minimal number of parameters. This way I shall study a minimal complex network model that can show persistence and as well as can shed light on the mechanism behind the persistence in the brain networks.

## 2 New and Noteworthy

There exists a debate in the Neuroscience on how the brain attains wakefulness. This may concern the state of the mind after some aetiology of coma, myoclonus, anaesthesia, etc. My work presents it as a phase transition of the network activity due to the action of inhibitory synapses. This claim has been further buttressed by using the Wilson Cowan Model. Then, it has been shown that the intrinsic currents through the ion channels give vent to the persistent activity, that in the absence of inhibition takes the form of oscillating global network activity. Its significance concerns myoclonic seizures in the hippocampus or other brain regions with abundant pyramidal neurons.

## 3 Model Formulation

### 3.1 FitzHugh Nagumo Network Model

The use of computational models to study neurons began in 1952 with the conductance-based Hodgkin and Huxley model, which explained the dynamics of the squid giant axon. One simplified version was presented by Dr Richard FitzHugh in 1961 and independently by Jinichi Nagumo *et al*. in 1962 [4],[5],[6]. FitzHugh Nagumo Model (FHN model or simply FHN) treats an isolated neuron as a slow fast dynamical system. According to the FHN, the neuron’s excitation potential (or voltage) is a dynamical variable with its differential exhibiting cubic nonlinearity. This excitation is modulated by another slow variable. For example, in the present paper, neuron voltage varies at a timescale 1*/*ϵ = 100 times faster than the modulation variable. Dynamically, this system can undergo supercritical and subcritical Andronov-Hopf bifurcations concerning the increasing stimulus, (see Figure 1). Figure 1 is alluding to codimension-one bifurcation [7],[8]. The system possesses canard (*duck-like*) solutions at stimulus valued from 0.111 mA to 0.11186 mA[9] Remarkably, the nonlinear dynamics of the model not only explain neuron behaviour but also elucidate dynamics of chemical oscillatory reactions, arrhythmia, naturally occurring spatiotemporal patterns, etc. (see e.g., [10], [11]) Furthermore, successful simulations of logic gates have been achieved using the model’s ability to create and switch between robust spatiotemporal patterns, underscoring its potential role in understanding the dynamics of the computation [12]. The mathematically synaptically connected network system is written as equations (2), (3) and (4).

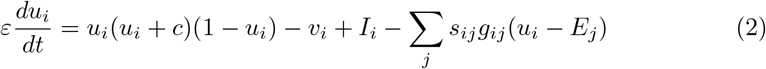

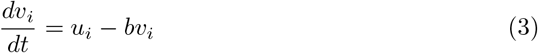

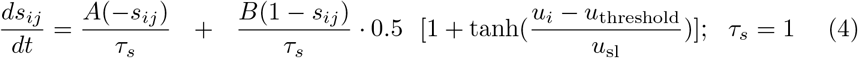

**Fig. 1.**
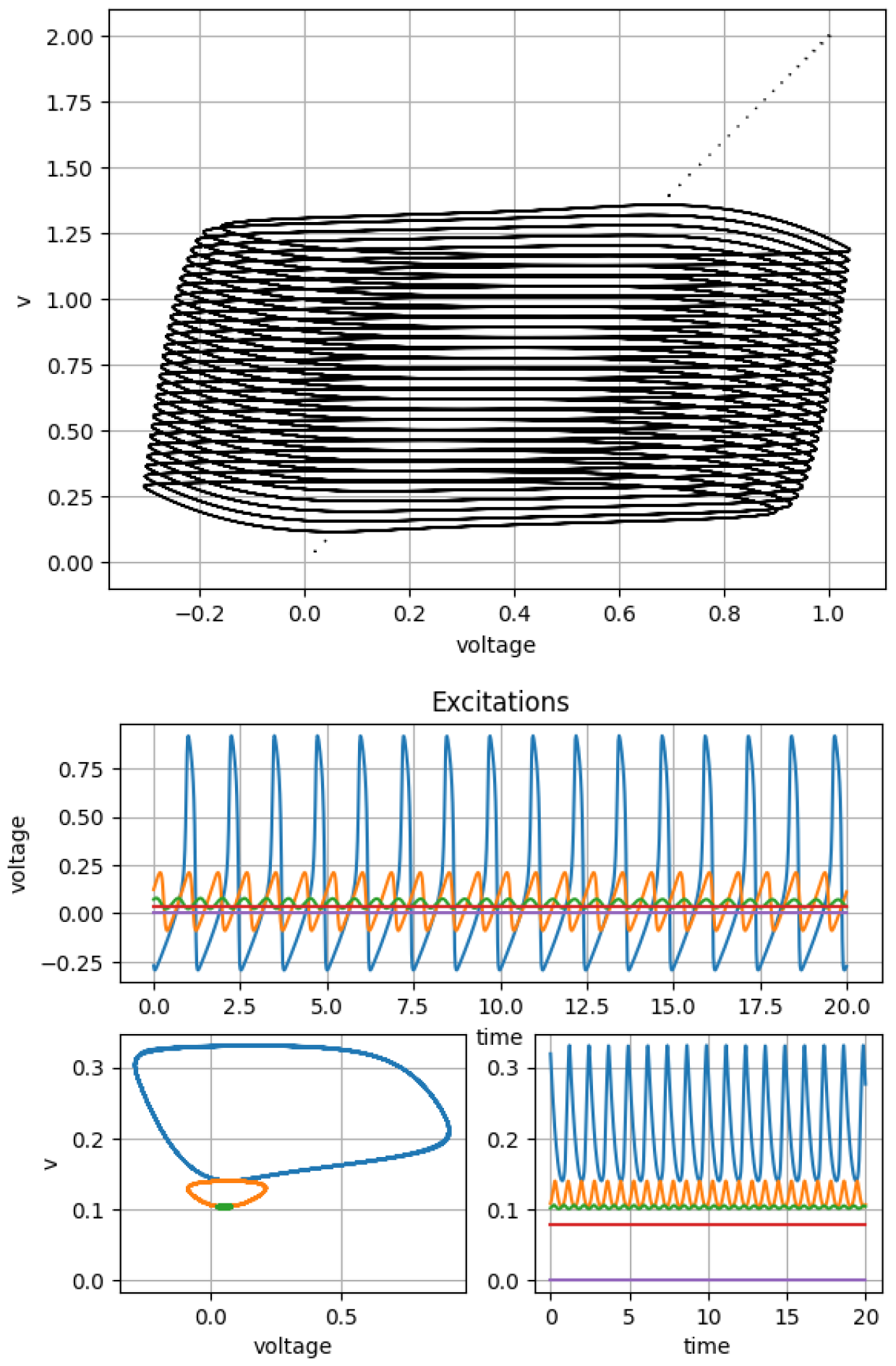
In the depicted phase space, the horizontal axis denotes the excitation (*u*) of the neuron, while the vertical axis represents the slow modulating variable (*v*). The stable dynamical state of the neuron within this phase space ascends with the incremental rise of current (*I*) from null to 2 mA, traversing initially through the supercritical and subsequently the subcritical Hopf bifurcation regimes. Noteworthy is the absence of persistent response to transient stimuli. The lower panel shows the supercritical Hopf Bifurcation.

**Fig. 2.**
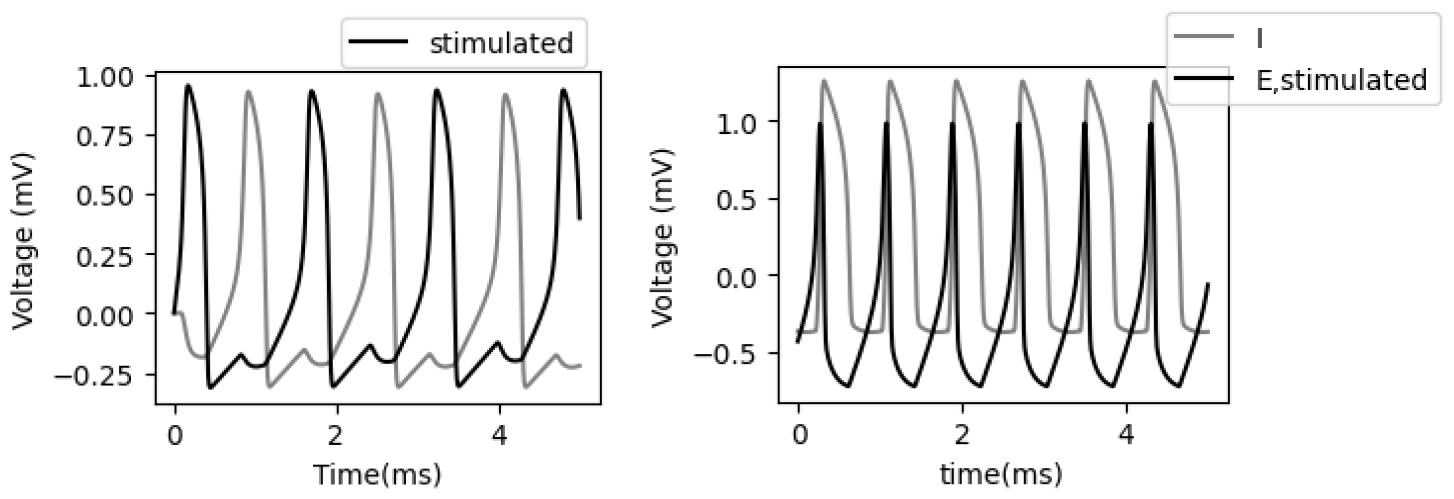
Left. Both the neurons are Inhibitory and bidirectionally synaptically connected with g = 0.05 units. Right. Stimulated neuron was changed to Excitatory and *g* was changed to 0.9 units. Black neuron was stimulated for 0.2 ms. For both cases, Sustained oscillations in response to a transient stimulus can be observed.

#### Parameters

- *u*_*i*_ and *u*_*j*_ are the presynaptic and postsynaptic neural potentials, respectively.
- *v*_*i*_ and *v*_*j*_ are slow modulation variables for presynaptic and postsynaptic neurons, respectively.
- The parameters *c* = −0.1 and *b* = 0.5 describe the model kinetics, and ϵ = 0.01, except in Figure 3 where it is 0.075, is the variable controlling the time scale differences between slow and fast variables. These values are chosen such that each uncoupled neuron is an excitable system.
- *I*_*i*_ is the external current or stimulus and varies throughout the study.
- *s*_*ji*_ denotes the synaptic gating variable.
- *g*_*ij*_ represents the entries of the synaptic weight matrix (or the adjacency matrix).
- *E*_*syn*_ represents the synaptic reversal potential, with *E*_*syn*_ = 5 for excitatory and *E*_*syn*_ = −5 for inhibitory neurons, respectively.
- Here, *A* = 3 and *B* = 3 are the rate constants, and *u*_*th*_ = 0.3 and *u*_*sl*_ = 0.001 are the parameters determining the shape of the synaptic term. The equation for the gating variable *s*_*ij*_ depends only on the membrane potential of the presynaptic neuron *V*_*j*_.
- 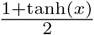 is a step-up function.

**Fig. 3.**
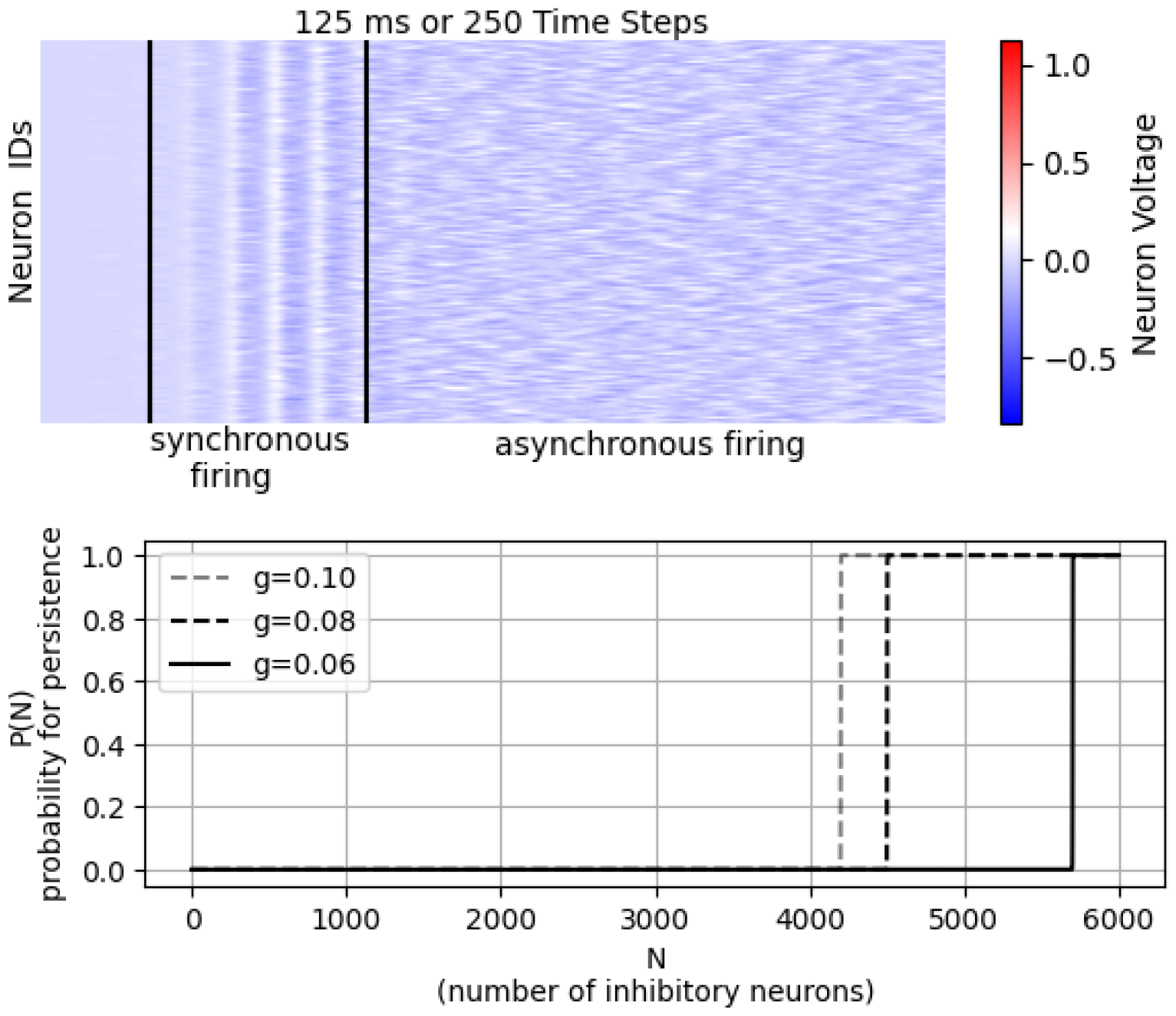
The P(N) on the Y axis is indicative of the probability of persistence concerning the number of inhibitory neurons N in the EI network with 6000 nodes. Hopf Bifurcation occurs on increasing inhibition in the network. Effect of synaptic conductance *g* is indicated. The top figure shows the spatiotemporal pattern for a neural persistent activity for 25 ms at N = 4800 and *g* = 0.10 The effect of initially being held by the inhibitory synapses on neural network activity and then being released from it can be seen as persistent neural network activity in *mV*.

The parameters that were varied throughout the work can be found in Table 1.

### 3.2 Firing Rate Model

Wilson Cowan Model equations describe the neural firing rates *F*_*E*_ and *F*_*I*_ of the interconnected excitatory and Inhibitory **populations** with weighted recurrent and lateral synapses that model the point stochastic process going on in the, say, spiking neural networks [13] using coupled ordinary differential equations with the form as equations (5) and (6) [14].

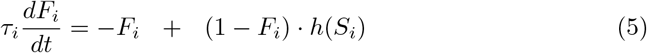

where

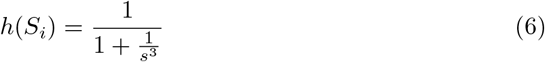

is the Firing Rate Response function. The specific functional form of *h*(*S*) varies in the literature. The present example is inspired by the recent work of Buxton [15]. If *S*_*E*_ and *S*_*I*_ are the synaptic currents entering excitatory and inhibitory populations, and *h* is the monotonically increasing firing rate response function ranging between zero and one; then, synaptic current entering into node *i* coming from node *j* is related to synaptic weights and biases of the concerned connection as in equation (7).

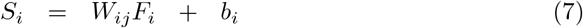

*S*_*i*_ can be positive or negative depending upon whether node j is excitatory or inhibitory. Throughout the work in this paper, Dale’s Law is true. *W*_1_, *W*_2_ physically means a measure for the number of excitatory synapses going into per excitatory and inhibitory neurons respectively and *W*_3_, *W*_4_ physically means a measure for the number of inhibitory synapses going into per excitatory and inhibitory neuron respectively [16]. This model has helped connect computational results to EEGs [14], [11], [17], [18], [19]. The list of the parameters used can be found in Table 2.

**Table 1.**
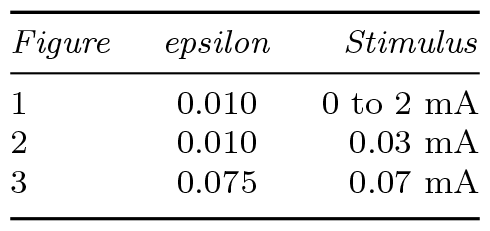
The FitzHugh Nagumo Parameters for Figures 1, 2, 3

| *Figure* | *epsilon* | *Stimulus* |
| --- | --- | --- |
| 1 | 0.010 | 0 to 2 mA |
| 2 | 0.010 | 0.03 mA |
| 3 | 0.075 | 0.07 mA |

**Table 2.** Wilson Cowan Model Parameters for Figure 4

| *Population* | *Parameters* | *Values* |
| --- | --- | --- |
| Crossed | $W_a, W_b, W_c$ | 0 to 2 mA |
| 1 | $W_1, W_2, W_3, W_4$ | 6.5,6.5,6.5,0 |
| 2 | $W_1, W_2, W_3, W_4$ | 6.5,6.5,6.5,0 |
| 3 | $W_1, W_2, W_3, W_4$ | 4,2,1.2,0 |
| 1 | $I$ | 0.32 for 0.5 s to $E_1$ |
| all | $\tau_{E1}, \tau_{E2}, \tau_{E3}$ | 0.010,0.010,0.010 s |
| all | $\tau_{I1}, \tau_{I2}, \tau_{I3}$ | 0.007,0.007,0.007 s |

**Table 3.** Prameter Values for Pinsky Rinzel Model for Figures 5 and 6 (1)

| $p$ | $g_c$ | $I_s$ | $I_d$ | $g_L$ | $g_{Na}$ | $g_{K-DR}$ | $g_{KC}$ | $g_{Ca}$ | $g_{K-AHP}$ | $g_{K-Ca}$ | $Cm$ |
| --- | --- | --- | --- | --- | --- | --- | --- | --- | --- | --- | --- |
| 0.5 | 2.1 | 0.5 | 0.0 | 0.1 | 30.0 | 15.0 | 15.0 | 10.0 | 0.8 | 15.0 | 3.0 |

**Table 4.** Prameter Values for Pinsky Rinzel Model for Figures 5 and 6 (2)

| $V_L$ | $V_{Na}$ | $V_K$ | $V_{Ca}$ | $V_{syn}$ |
| --- | --- | --- | --- | --- |
| 0 | 120 | -15.0 | 140 | 30 |

### 3.3 Pyramidal Neurons

Pyramidal Neuron is a common class of neurons found in the cerebral cortex of apparently every mammal, as well as in Avians, Pisces, and reptiles. Pyramidal neurons are also common in subcortical structures such as the hippocampus and the amygdala and they comprise about two-thirds of all neurons in the mammalian cerebral cortex. There are two dominant families of neurons in the cortex, excitatory neurons, which release the neurotransmitter glutamate, and inhibitory neurons, which release g-amino-butyric acid (GABA). Pyramidal neurons are the most populous members of the excitatory family in the brain areas they populate [20]. These were first modelled computationally by John Rinzel and Paul F. Pinsky in detail [21]. Their model divided the neuron into two interconnected neural compartments: Soma and Dendrite; with their respective excitations/voltages *V*_*s*_ and *V*_*d*_. Apart, their model included calcium current dynamics and the dynamics of other intrinsic ionic currents through modelling probabilistic gates for ion channels through the use of so-called *gating variables*. AMPA (*α*-amino-3-hydroxy-5-methyl-4-isoxazolepropionic acid) and NMDA (N-methyl-D-aspartate) are types of ionotropic glutamate receptors found at excitatory synapses in the central nervous system. AMPA synapses are fast-acting and primarily responsible for mediating fast excitatory neurotransmission. When glutamate binds to AMPA receptors, it triggers the influx of sodium ions into the postsynaptic neuron, leading to depolarization and excitatory effects. NMDA synapses, on the other hand, are involved in a variety of processes including synaptic plasticity, learning, and memory. Activation of NMDA receptors requires the binding of glutamate as well as the presence of a co-agonist, typically glycine. NMDA receptors are unique in that they are both ligand-gated and voltage-dependent. They allow the influx of calcium ions in addition to sodium ions, which can trigger various intracellular signalling cascades important for synaptic plasticity and long-term potentiation (LTP). Pinsky and Rinzel also included the dynamics of the AMPA and NMDA synapses and examined the network properties by modelling them as event-driven connections [21]. For equations and values used, refer to section (5): Methods.

## 4 Results

### 4.1 Dynamics of Single Neuron

One single FitzHugh Nagumo neuron can not show persistence in the face of an absence of external stimuli. Its solutions (*u, v*) will eventually touch the boundaries (here *u-nullcline*) and shall slide to zero. In the presence of different stimuli ranging from 0 to 2 mA, the stable states are shown in the 1. The figure also alludes to the ongoing supercritical and then subcritical Hopf on increasing stimulus *I* step-wise. More can be found on [9].

### 4.2 Dynamics of Coupled Neurons

A couple of nonlinear (relaxation) oscillators are different from a couple of linear oscillators because the superposition principle does not apply in the former system. Therefore, it can produce rich dynamical states that the latter cannot produce. It can switch between these dynamical states (or bifurcate in a variety of ways) passing through intermittent states when the model parameters change. Here these parameters can be *I*_*i*_, *E*_*i*_, *I*_*j*_, *E*_*j*_, *g*_*ij*_, *et cetera*. These also promote the synchronization of the two oscillators. Books, for example, [7], [8], [22], [23], [24], etc. provide ample information on this subject, and thus, preempting the need for repetition. So, this study delves directly into the discussion of whether two recurrently coupled nodes with synaptic connections can sustain activity through self-driving each other when transiently stimulated. It is well-known that the activity patterns of coupled neurons vary depending on (1) the type of synapses/neurons, i.e., excitatory or inhibitory, and (2) the coupling strengths [9]. Through numerical simulations, I found that in the case of an inhibitory presynaptic neuron, the postsynaptic neuron fires an action potential after being released from the suppressing effect of the inhibitory presynaptic neuron. This rebound effect is a factor that I found responsible for the persisting neural oscillation in the system of synaptically coupled FitzHugh-Nagumo neurons, see for example, 2. In literature, this is called as the post-inhibitory rebound, e.g., see [8] on page 242. Thus, if both the synapses (or equivalently neurons here) are excitatory, activity cannot be sustained. Furthermore, strong synaptic conductance can convert this rebound to a sustained activity pattern 2. In the case of EI or equivalently *IE* pairs, the strength of the synaptic conductance plays a key role in maintaining sustained oscillations. If the inhibitory synapses are fast, i.e., *τ*_*s*_ in equation (4) is increased, it shall cause the suppression long-lasting, thus, affecting the frequency of the oscillations, if they exist at all. Previous analyses of two mutually coupled neurons, or networks operating in regular firing mode, have also shown that the oscillatory properties, particularly the frequency of the global oscillation, depend strongly on the decay time of inhibitory postsynaptic currents (see, for example, [25] [26],[27],[28]). Therefore, the results and reasoning of the present study agree with the conclusions of these landmark studies.

### 4.3 Sparsely Connected EI Network

For the study of the brain, the realistic form of sparseness is the one in which a synapse is present in the model network with probability C/N at each pair of cells, independently from pair to pair. This is a claim found corroborated by [13]. Biologically as well, the number of arriving synaptic inputs varies from neuron to neuron and makes up a probabilistic distribution. More on sparseness can be found in [29]. Secondly, since the neuronal networks are random, their properties can be studied with the help of Erdos Renyi’s random graphs. As the number of nodes in the net tends to achieve infinity, i.e., assuming the thermodynamic limit for the number of nodes in the graph, then the probability of having a specific type of edge, say, the one that represents self-connection; is vanishingly small or zero [30]. Also because of the experimentally observed absence of self-connections in real neuronal networks, we do not model self-connections. My study found a supercritical Hopf bifurcation achieved through decreasing the **Excitation Inhibition ratio** that develops a separation between the network activity regions with stationary and oscillatory global activity (3). This finding of oscillatory activity in networks of mostly as well as purely inhibitory neurons agrees with previous modelling studies with either simpler or detailed conductance-based neural models, as in, the modelling of the Gamma Oscillations by Wang Buzsaki in 1996. [31]. Others like [25], [32], [33],[13] and with evidence from both *in vivo* [34] and *in vitro* [35], [32] and [36] experiments corroborate that inhibition plays an important role in the generation of network oscillations. For Figure 3, (a) five randomly chosen nodes in the network were provided stimulus valued at 0.07 mA for 1 ms. The persistent activity is shown in the figure for 25 ms but it continues beyond (b) among network theory measures, only the study of the percolation process on a random graph (called directed percolation) alongside its entailing First Order Phase Transition is conducted [37]; as we vary *N*, i.e., the number of inhibitors while keeping total population fixed. The neural network activity seen in the middle (marked as synchronous firing) of the 3 is the Burst Suppression (SB) electroencephalogram (EEG) pattern and is a hallmark of a brain that is not fully awake either due to it being anaesthetized (e.g., say through the use of parenteral barbiturates) or in a coma. I ruled out myoclonus as it occurs for lack of inhibition, not the contrary [38] [16] [39]. It may also be the Gamma Oscillations in the network of Inhibitory neurons as found previously [31].

The third pattern in 3 models the *spontaneous brain activity* of a healthy functioning brain [40]. Either way, all in all, physically this transition marks up the *awakening* of the brain. It has been caused due to the presence of inhibitory synapses. This finding is consistent with the prior findings that the inhibitory neurons does cause oscillatory activity in the brain. Also, a recent model for the study of anaesthesia has claimed a role of inhibition and a self-organised threshold for it in causing anaesthesia [16]. My study is significantly new, as it proposes that the wakefulness from the anaesthesia is also caused by the inhibitory synapses. To understand the effect of GABA synapses with delays, the reader is directed to start at the reference [31] and [41].

### 4.4 Firing Rate Model

The spatiotemporal pattern shown in Figure 3 is subtle. Network activity undergoes three phases: No Firing, Synchronized Firing, and Asynchronous Firing all by itself through some sort of self-organized criticality (SOC) concerning the inhibition present in the network. To break this down; I proceed as follows: three effects that can be observed in the spatiotemporal pattern are, One, no activity for the first few milliseconds (1-80 ms) if the stimulus existed for the first 1 millisecond, Two, self-initiation of the activity in the form of synchronized neural firing (or equivalently oscillating firing rate, i.e., number of neurons firing per millisecond) that oscillates near zero and continuing such for nearly 100 ms, and Three, transitioning of this oscillating firing rate to a near stationary firing rate or equivalently asynchronous neural firing which is stable/saturated at some higher firing rate value. For definitions see [13] and [42]. I assert that the observed 2 phase transitions are caused by inhibition (or inhibitory synapses) only, instead of any other network measure like structure, finite size effect, complexity, *et cetera*. Further, while testing for this hypothesis in this subsection, I also impose that my theory must be useful for understanding the neural activity caused by the neurons in the brain. This all can be readily achieved using the Wilson Cowan model as a two-edged sword as described below. The goal of using the Firing Rate (also known as, mean-field) model presently is to test the observation that the synaptic strengths alone are driving the network towards persistence in the way observed in the previous subsection. All in all, this exercise will help us understand (a) why activity remains suppressed initially (b) why synchrony (c) why transition to asynchronous firing from synchronized firing. On the second hand, the firing rate model helps to establish a connection among the figure 3 and the EEG patterns observed clinically say during, say anaesthesia, sleep, coma, (no seizure), spontaneous brain activity *et cetera*. The results obtained are shown in 4. As we are considering a random network, the possibility of the structure having anything to do with this is ruled out. Figure 4 shows the circuits of the neural populations I considered to recreate the observations. As visible, I considered three interacting Wilson Cowan populations, a scheme historically under application since the 1980s, see for example, [1]. As per the circuit diagrams shown in Figure 4, the third EI population (*E*_3_*I*_3_) can inhibit the first excitatory population (*E*_1_) through either exciting its locally coupled inhibitory population *I*_1_ or directly inhibiting it using *I*_3_ by establishing the connection specified by synaptic weight *W*_*c*_ and zero bias. Such connections mimic long-range connections in the brain. Since the inhibitory population can’t make long-range connections, only the former of the two cases is biophysical. Rest, the inhibitory population is locally coupled to the self-feeding excitatory population. Excitatory populations connect among themselves and these connections mimic short-range connections of the brain. This configuration of three WC EI nodes can produce a wide variety of persistent dynamical patterns that are also observed in the EEG. As can be seen in 4, the doubling of the weight *W*_*c*_ of the inhibitory synapse from 2 to 4 causes the **sustained** firing of the neurons to begin; provided the “duration of the applied-current” crosses a certain threshold (here, 400 ms). For a neural network in which inhibition dominates, when the inhibitory synaptic current flows for a duration that is either equal or greater than a threshold value (e.g., here 400 ms), the synchronized firing phase of the network global activity transitions to the asynchronous firing state. The reason is the spontaneous symmetry breaking by the neurons/constituents of the network [43].

**Fig. 4.**
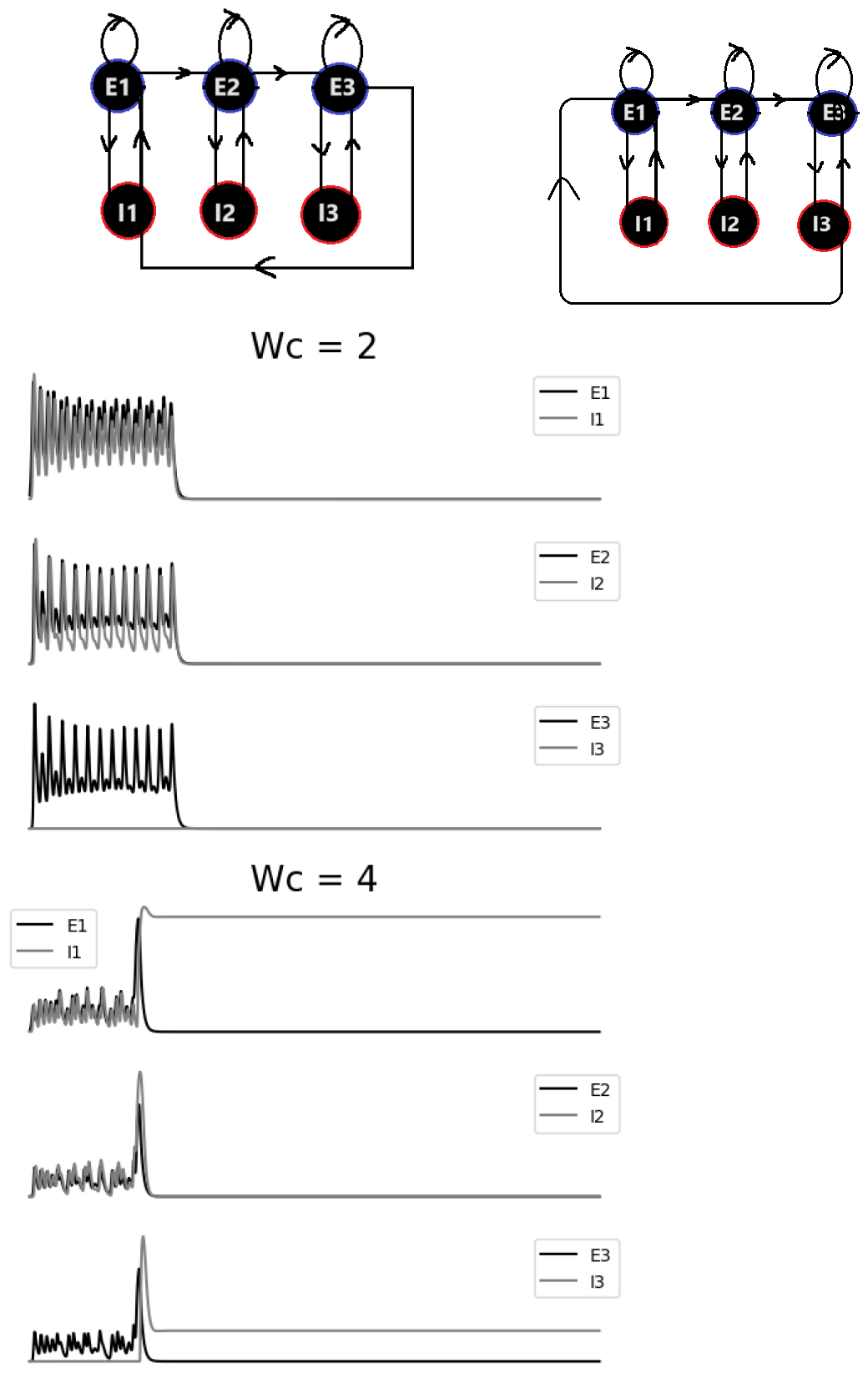
**Firing Rate Model**. Stimulus was provided to *E*_1_ for 0.5 seconds. The x-axis shows a time interval spanning 2 seconds. The y-axis is the Firing Rate. In the top two figures, two possible circuits with three interconnected Wilson Cowan Populations are drawn. Below, it is shown that if the inhibition in the network is not strong enough, as measured by the synaptic weight of the longrange inhibitory synapse *W*_*c*_, neural persistence can not be achieved. However, if it is increased to 4, giving a model of densely connected neurons with strong inhibition, neural persistence is achieved, provided, the applied current duration lasts for at least 400 ms. After 400 ms of the strong inhibition, a phase transition happens causing the network to switch from the oscillatory to a stationary global firing rate state. The reason is that the spontaneous symmetry breaking in the network results in the desynchronization of the neurons “all at once” thus resulting in the apparent phase transition.

### 4.5 Network of Pyramidal Neurons

To preempt the possibility of the propagation of the claim through my work that the neural network models constituted by the excitatory nodes and synapses can not have persistence as their dynamical state, I present my results on this subject using a detailed neural network model constituted by the Pinsky Rinzel (pyramidal) Neurons [21] with separate Soma and Dendrite compartments and probabilistic ion channels, sparsely and randomly connected by two different synapses AMPA and NMDA. Figure 5 right panel shows the result of stimulating 40 out of 1000 densely connected PR neurons of a homogeneous network for 1 ms. The resulting persistence is shown for 50 ms in right of the Figure 5. This persistence has been seen to go on for at least 500 ms. The result of the interaction among soma and dendrite, many intrinsic currents, and excitatory synaptic currents in the network gives neural persistence after 500 ms. The global oscillatory frequency has been seen to increase, as the *g*_*AMPA*_ strength is manually increased from 0.005 to 0.01 to 0.015 to 0.02 keeping *NMDA* synapse non-functional, which is completely in favour of the work conducted by Paul Pinsky and John Rinzel themselves [21]; though in their work, the persistence as a possible dynamical was not studied. In conclusion, though the network system constituted all or most of the excitatory constituents can not show persistence but equipped with synaptic gates modulating currents, as in the Pinsky Rinzel network model, the system expands its dynamical phase space to a large number of dimensions and hence finds the possibility for persistence [44]. In Figure 6, the persistent spatiotemporal pattern (or semiflow) characterized by the hypersynchronous activity of the pyramidal neurons in the absence of inhibition is symbolic of epilepsy; a neurophysiological disease caused due to excessive neuronal excitation due to either absence of or low lying inhibitory firing rate. It is particularly common in brain regions containing many interconnected pyramidal neurons, such as the hippocampus [20]. The activity pattern seen is called the Suppression Burst pattern; a consequence of disruption in EI Homeostasis and characterize, e.g., neonatal myoclonus, etc. on EEG [38]. Due to the ability of the network of pyramidal neurons to show robust persistence, they also play a profound role in memory formation [45].

**Fig. 5.**
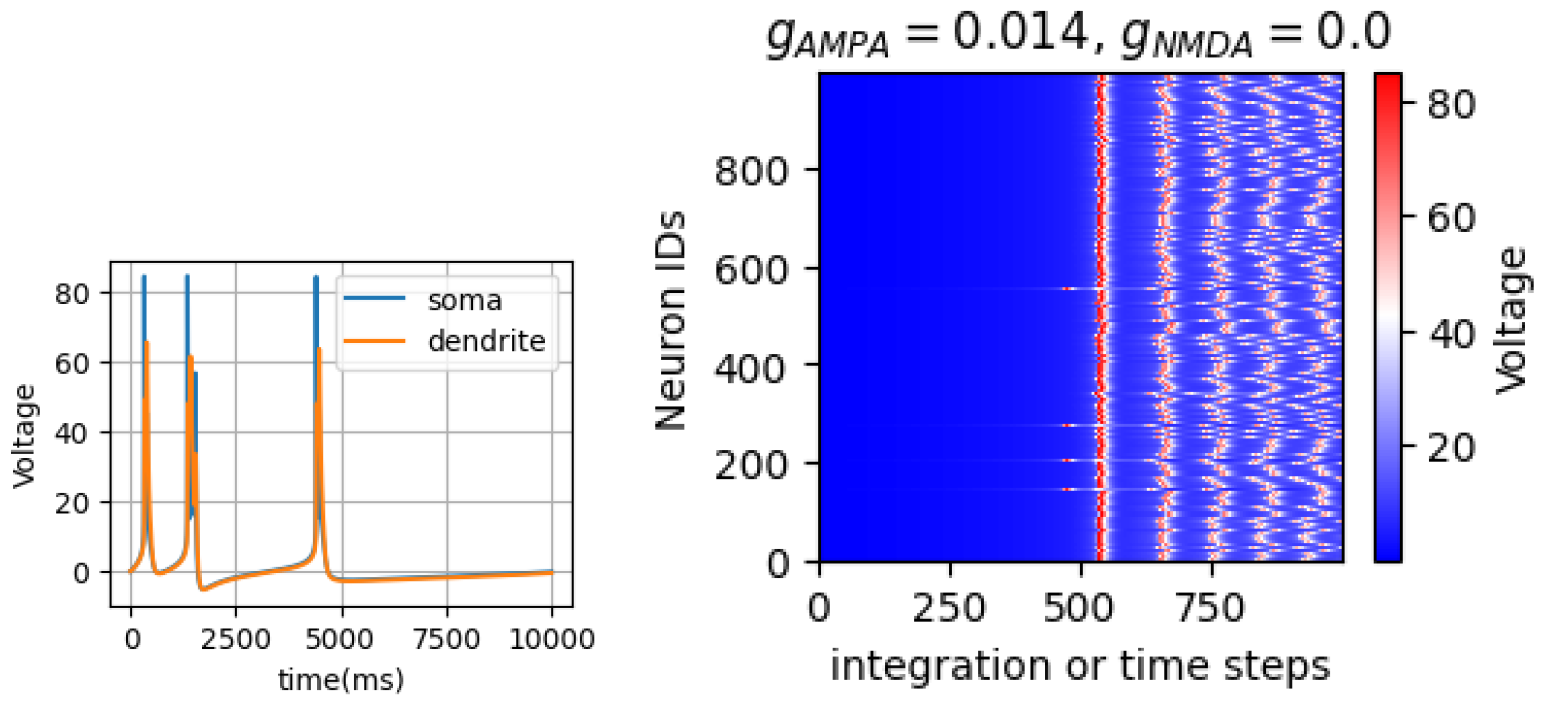
Left. Single Pinsky Rinzel Neuron. *V*_*s*_ is somatic voltage (mV) and *V*_*d*_ is dendritic voltage (mV). 0.5 mA current was provided consistently. Right. A network of 1000 such neurons was stimulated for 50 ms. The current of 0.5 mA was provided to 40 random neurons for 1 ms. Oscillating collective neural voltage (in mV) was observed.

**Fig. 6.**
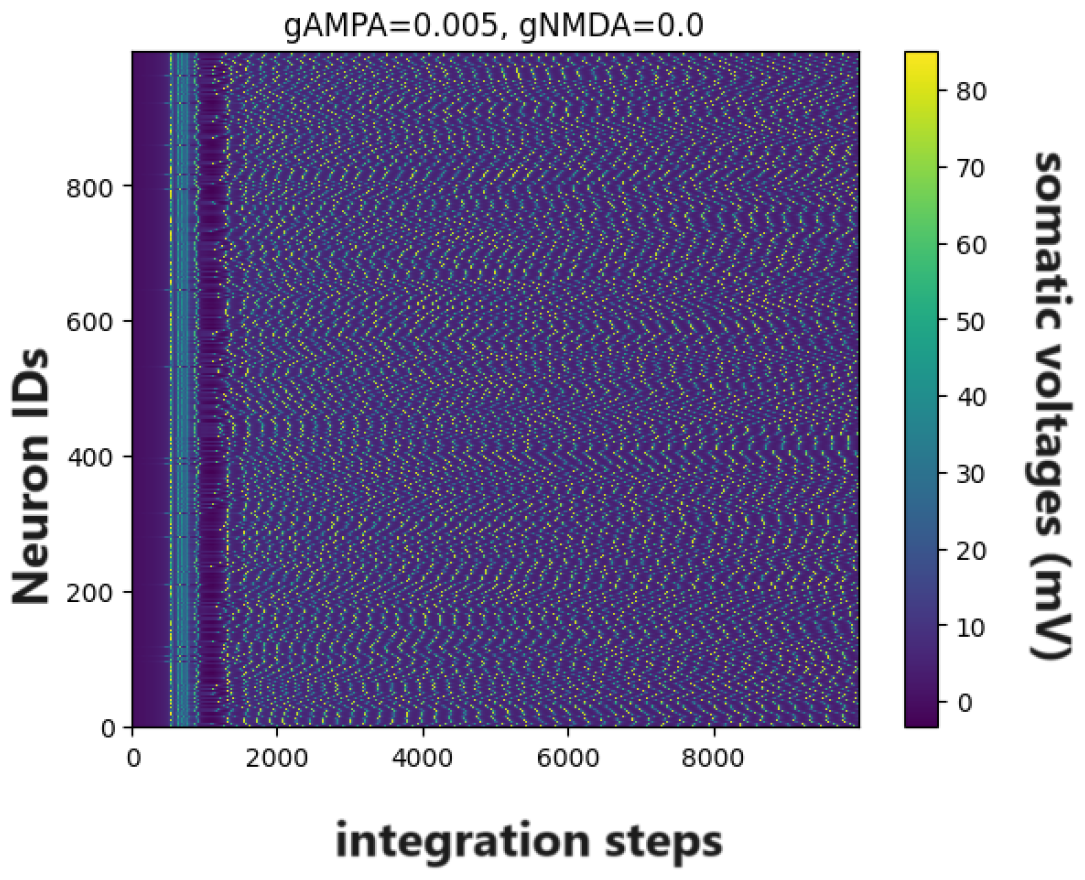
Pinsky Rinzel Network Model. Current supplied to 40 (out of 1000) randomly chosen neurons for 1 ms. Simulation time is shown for 500 ms. NMDA is not present. The major difference between the two is that the NMDA-type receptors (NMDARs) typically possess slower kinetics than AMPAtype receptors (AMPARs)[50]. Semiflow can be seen above.

### 4.6 Nonlinear Dynamics

Figure 3 shows a **semiflow** defining the spatiotemporal evolution of the Fitzhugh Nagumo network on a subset of its state space, say *X*, a subset of a set consisting of discrete time steps *J* and defined by a map (or function) Φ such that; Φ maps either a subset of or entire *J* × *X* → *X* to *X* while also satisfying the semiflow property, see equation (8) and section 7].

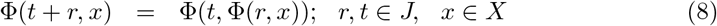

I ask whether the dynamical system persists as a whole or at least in parts when the fraction of one of the subpopulations rises (denoted by N). This question can be formulated by using a *persistence function* (here, neuron voltage as shown in the colorbars) such that it maps state space to the ℝ^+^. This way the FHN system that I considered is called N-persistent. This can be seen in Figure 3. There exists a well-worked-out theory aimed at classifying the persistent dynamical states as Uniform, Strong, Weakly Uniform, et cetera [3]. In our case, it won’t be of much help. The activity of the individual neuron in the network remains bounded by the lower and upper bounds which are invariant with respect to the passing time. In the present case, it is rather the question of why some neurons in the sparse random network started firing sans synchronization suddenly, though persisted the way they were persisting exactly before. Neural networks such as FHN and PR started with some degree of symmetry, where all nodes have similar connectivity patterns or attributes. As interactions and connections evolved within the network, small fluctuations or perturbations arose due to various factors like node preferences, external influences, or random events. These fluctuations amplified over time, leading to the emergence of hubs or clusters within the network. Some nodes may have acquired more connections or influence than others, breaking the initial symmetry. Once certain nodes or clusters gained prominence, they attracted more connections or resources, further reinforcing their dominance within the network. At a critical point, the network undergoes a phase transition where the symmetry is spontaneously broken, and a new organizational structure emerges. This transition can lead to the formation of distinct communities, hierarchical structures, or functional specialization within the network. Spontaneous symmetry breaking in neural complex networks often leads to self-organization, where the network’s structure evolves towards states of higher efficiency, resilience, or adaptability. Overall, spontaneous symmetry breaking in my case occured through the interplay of network dynamics, amplification of fluctuations, and the emergence of dominant nodes or clusters. Understanding these processes can provide insights into the organization and behavior of information flow, and neuropathologies in brain networks.

## 5 Methods

### 5.1 General

The number of nodes for the FHN network and PR network are 6000 and 1000 respectively. Both networks are sparse and random satisfying that each neuron has the probability of having a link being 0.006. In other words, the neurons in the network have an average degree of 6, a type of dense homogeneous network prominent in network science (see for example, [37] [29] for more information on random nets). Further, the models FHN, WC and PR were numerically integrated using the Runge Kutta 4 method using step sizes of 0.01, 0.001 and 0.05 respectively in *Python*. The parameters that were varied in The Fitzhugh Nagumo Model are given in Table 1. Next, the parameters that were varied in The Wilson Cowan Model are given in Table 2. Further, in Wilson Cowan model, *W*_*a*_, *W*_*b*_, *W*_*c*_ are the synaptic weights of the connections that connect *E*_1_ to *E*_2_, *E*_2_ to *E*_3_ and *E*_3_ to *I*_1_ (or equivalently *I*_3_ to *E*_1_) respectively. *W*_1_, *W*_2_, *W*_3_, *W*_4_ are the synaptic weights of the connections that carry current from *E*_1_ to itself, *I*_1_ to *E*_1_, *E*_1_ to *I*_1_ and *I*_1_ to *I*_1_ for anyone WC node (see Figure 4 top panel). Biases are zero, except for the synaptic connection that connects *E*_1_ to *E*_1_, which is 0.32 for the first half second and otherwise zero. The inhibitory population is faster than the Excitatory population, as in, the brain’s Interneurons are faster than the Pyramidal Neurons. The values of the parameters used in Figures 5 and 6 are as in tables for PR Network 3 and 4.

### 5.2 Pyramidal Neuron Model

Equations (9) – (31) define the Pinsky Rinzel Model.

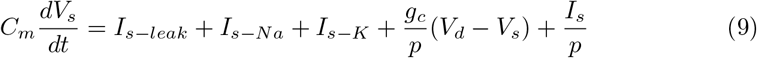

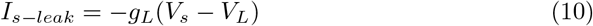

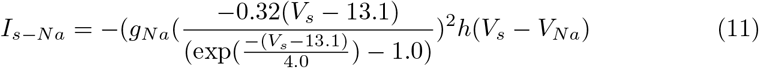

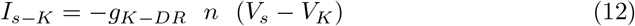

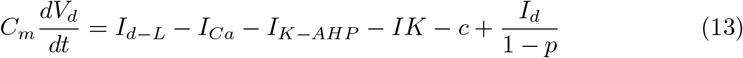

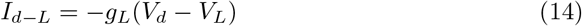

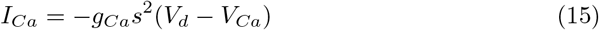

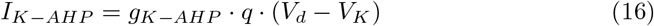

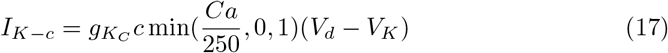

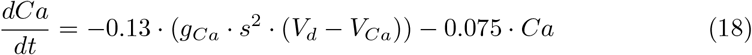

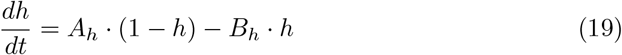

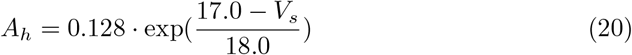

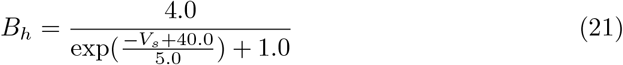

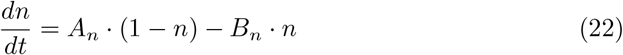

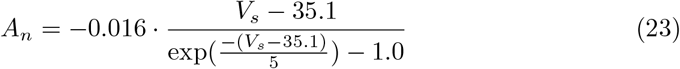

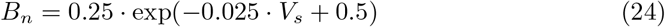

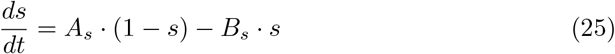

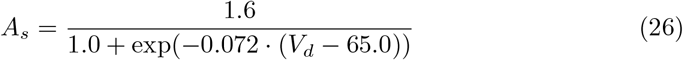

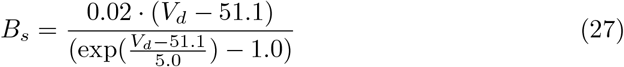

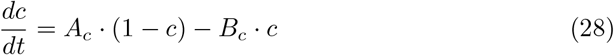

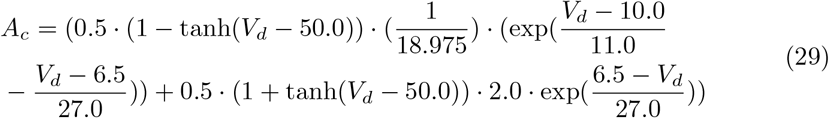

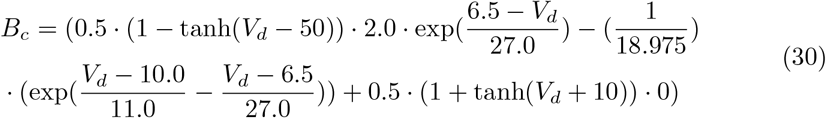

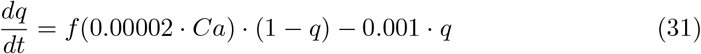

## 6 Discussion

The results suggest that persistent neural activity stems from inhibitory synaptic actions, making it susceptible to the influence of transmission delays and the probabilistic distribution of synaptic time scales inherent in the brain. These factors can induce synchronization within the network, leading to regularities in neuronal firing patterns and subsequently impacting global activity to either its advantage or otherwise. This shall be equivalent to tuning two synaptic time scales and the average degree in the neural network model. This hypothesis finds support in previous modelling experiments such as [25], [26], [27], [28], [45], etc. Recent work by Christopher M. Kim *et al*., building upon the approach of Nicolas Brunel using an EI population model, significantly addresses this aspect, thus obviating the need for repetition here [42],[13]. Memory, defined as enduring alterations in system states in response to environmental cues, enables the retention of information about transient signals long after exposure [46]. Nicolas Brunel in the brain and more recently, Mitra *et al*., in cellular signalling, have directly linked persistent activity to memory functions [13][47]. In addition, this model interfaces intricately with the realm of analog computing, an arena currently undergoing a notable resurgence of industrial fascination [48], [49]. Therefore, it shall also be interesting to incorporate the energy consumption constraint on the neural network model and observe its activity; a practice that may open gates to understanding the emergence of the seizures (e.g., infantile myoclonic epileptic encephalopathies) as well [38],[16] and [15].

## 7 Conclusion

I discussed the percolation effect on a sparse and random network of FitzHugh Nagumo excitatory neurons by increasing the fraction of inhibitory neurons in the population and observed an adiabatic phase transition from dead to persistent network activity 3. This happened due to the spontaneous symmetry-breaking effect by neurons in the network, when the network is (a) dense and (b) inhibitory enough [43]. Other types of percolation effects concerning synapses are possible to give significant results (like synaptic delays, average degree distribution, etc.). This whole constituted ***synaptic*** cause for persistence. I also saw that the ion channels in a neuron may lead to persistence. This constitutes ***intrinsic*** cause for persistence and is seen in all excitatory neural networks. I conclude, based on numerical evidence, that wakefulness can be attributed to the presence of inhibitory constituents in the brain. Memory can be attributed to the intrinsic currents along with synaptic currents. Then, the role that can be played by the Nonlinear Dynamics to find out the necessary mathematical constraints on the neural network system is highlighted in subsection 4.6. This shall be elaborated on in future studies by introducing the subject of spontaneous symmetry breaking in the neural networks.

## Declarations

### Ethics approval and consent to participate

Not applicable

### Consent for publication

The author consents for publication.

### Data availability

Not applicable.

### Materials availability

Not applicable

### Code availability

Code is available on the internet on [GitHub](https://github.com/im-vishalverma/neural_persistence).

## Author contribution

Verma, V. performed the experiments and wrote the manuscript.

## A. Semiflow Property

In the realm of dynamical systems theory, the concept of a “semiflow” stands as a profound testament to the intricate interplay between time and evolution. A semiflow encapsulates the essence of continuity and evolution in systems where change is not merely a discrete event but an ongoing, continuous process.

At its core, the semiflow property embodies the idea of temporal coherence: the trajectory of a system’s evolution remains consistent and smooth over time, adhering to the laws that govern its dynamics. Formally, a semiflow is defined as a family of mappings, each representing the evolution of the system at different points in time, with the crucial stipulation that these mappings preserve both the temporal order and the continuity of trajectories.

In mathematical terms, let *X* be a topological space representing the state space of a dynamical system. A semiflow on *X* is a continuous mapping Φ : ℝ_+_ × *X* → *X*, where R_+_ denotes the non-negative real numbers, satisfying the following properties:

1. **Initial Condition Consistency**: The mapping Φ preserves the initial conditions, ensuring that for each *x* in *X*, Φ(0, *x*) = *x*. This foundational property anchors the evolution of the system to its starting point, providing a coherent basis for subsequent dynamics.
2. **Temporal Continuity**: The semiflow Φ exhibits temporal continuity, meaning that for any *t* ≥ 0, the mapping Φ(*t*, ·) is continuous. This continuity ensures the smoothness of trajectories as time progresses, highlighting the inherent stability and coherence of the system’s evolution.
3. **Semigroup Property**: The mappings Φ(*t* + *s*, ·) and Φ(*t*, Φ(*s*, ·)) coincide for all *t, s* ≥ 0. This property encapsulates the essence of time-invariance within the system, affirming that the evolution of the system over successive time intervals remains consistent, irrespective of the starting point.

The significance of the semiflow property reverberates across various domains, from mathematical dynamical systems to the modelling of physical phenomena. Its elegance lies in its ability to capture the essence of continuous evolution, providing a powerful framework for understanding the temporal dynamics inherent in complex systems. Indeed, the semiflow property stands as a testament to the profound synergy between time and evolution, illuminating the intricate tapestry of dynamics that governs the behaviour of systems in motion. As we delve deeper into the realm of dynamical systems theory, the semiflow property remains a beacon of insight, guiding our understanding of the timeless dance between continuity and change. Refer to Figure 7.

**Fig. 7.**
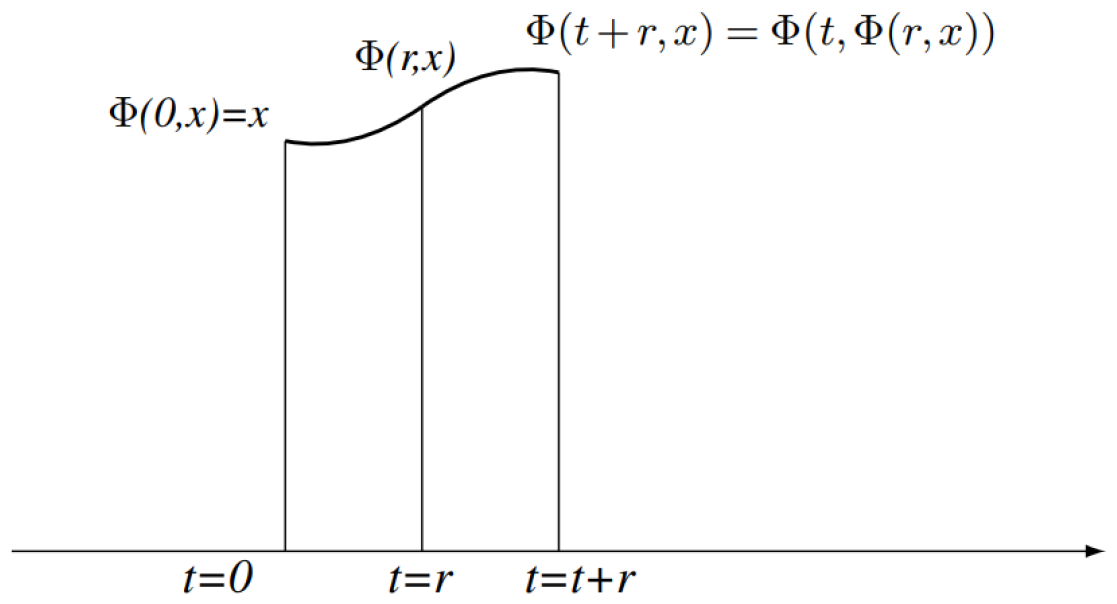
Semiflow property or Chapman-Kolmogorov equation. Φ(*x, t*) is sometimes itself called as the semiflow.

